# The PP2A-like phosphatase Ppg1 mediates assembly of the Far complex to balance gluconeogenic outputs and adapt to glucose depletion

**DOI:** 10.1101/2023.07.03.547494

**Authors:** Shreyas Niphadkar, Lavanya Karinje, Sunil Laxman

## Abstract

To sustain growth in changing nutrient conditions, cells reorganize outputs of metabolic networks and appropriately reallocate resources. Signaling by reversible protein phosphorylation can control such metabolic adaptations. In contrast to kinases, the functions of phosphatases that enable metabolic adaptation as glucose depletes are poorly studied. Using a *Saccharomyces cerevisiae* deletion screen, we identified the requirement of PP2A-like phosphatase Ppg1 for appropriate carbon allocations towards gluconeogenic outputs – trehalose, glycogen, UDP-glucose, UDP-GlcNAc – specifically after glucose depletion. This homeostatic Ppg1 function is mediated via regulation of the assembly of the Far complex - a multi-subunit complex that tethers to the ER and mitochondrial outer membranes as localized signaling hubs. We show that the Far complex assembly is Ppg1 catalytic activity-dependent. The assembled Far complex is required to maintain gluconeogenic outputs after glucose depletion. Glucose in turn regulates Far complex abundance. This Ppg1-mediated Far complex assembly, and dependent control of gluconeogenic outputs enhances adaptive growth under glucose depletion. Our study illustrates how protein dephosphorylation is required for the assembly of a multi-protein scaffold present in localized cytosolic pools, to thereby enable cells metabolically adapt to nutrient fluctuations.

## Introduction

Cells continuously adapt to fluctuating nutrient conditions by rewiring their metabolism to efficiently utilize available nutrients. Cells utilize multiple signaling systems for this metabolic rewiring, thereby facilitating adaptation to these environments (Broach, 2012; Efeyan *et al*, 2015; Lindsley & Rutter, 2004; Zaman *et al*, 2008; Smets *et al*, 2010). Therefore, understanding how signaling processes respond to changing conditions and regulate metabolism is of obvious interest. We have a growing understanding of signaling mechanisms that are active in specific nutrient environments, but our understanding of their roles in changing nutrient environments remains incomplete. However, this understanding is important to comprehend the basic principles of metabolic adaptation, with useful applications in industrial uses.

*Saccharomyces cerevisiae* is a remarkable model system using which several molecular mechanisms of metabolic adaptation, and conserved signaling systems that regulate these have been discovered (Zaman *et al*, 2008; Conrad *et al*, 2014). Particularly, this system has been instrumental in advancing our understanding of mechanisms of how cells adapt as glucose levels change. In glucose-replete conditions, *S. cerevisiae* cells preferentially utilize glucose as a carbon source and repress the utilization of other carbon sources, as well as mitochondrial respiration (Rolland *et al*, 2002; Deken, 1966; Gancedo, 1998). This includes repression of metabolic processes utilizing alternative carbon sources such as gluconeogenesis, glyoxylate cycle, TCA cycle, etc (Carlson, 1999). When starting in glucose replete conditions, *S. cerevisiae* cells adapt over the course of growth as they rapidly consume and deplete glucose (Käppeli, 1987). As glucose depletes, *S. cerevisiae* cells undergo a diauxic shift where cells reorganize their metabolic networks to utilize other available carbon sources (Schüller, 2003; Murphy *et al*, 2015; Brauer *et al*, 2006). In the post-diauxic phase, cells increase mitochondrial respiration and initiate gluconeogenesis to produce biomass precursors for growth. The outputs of gluconeogenesis include trehalose, glycogen, UDP-glucose, UDP N-acetyl glucosamine and others, all of which have essential roles in growth and adaptation (François & Parrou, 2001; Varahan *et al*, 2019a). While the roles of several signaling proteins in regulating carbon metabolism in high or low glucose respectively have been studied (Kim *et al*, 2013; Ljungdahl & Daignan-Fornier, 2012), our understanding of signaling mechanisms that regulate metabolic adaptations when glucose depletes remains incomplete.

Signaling mechanisms involve signal relays regulated by post-translational modifications (PTMs), particularly reversible protein phosphorylation, and can regulate metabolism (Oliveira *et al*, 2012; Humphrey *et al*, 2015; Wilson & Roach, 2002; Tripodi *et al*, 2015). In such signal relays, there typically is reciprocal regulation by writer-eraser systems, such as kinases and phosphatases (Jin & Pawson, 2012). In the context of glucose responses studied in yeast, our current understanding is kinase centric. In glucose-replete conditions, we have a growing understanding of regulation mediated by protein kinases such as Protein kinase A (PKA) and TOR kinase (Target of rapamycin), which activate growth programs in glucose-replete conditions. Specifically, these kinases activate transcription programs that control carbon metabolism, ribosome biogenesis, stress response, etc (Kunkel *et al*, 2019; Plank, 2022; Zaman *et al*, 2008; Thevelein & Winde, 1999; Galdieri *et al*, 2010; Pedruzzi *et al*, 2000; González & Hall, 2017). Upon glucose depletion, the Snf1 kinase (AMP-activated protein kinase) is activated and drives the utilization of other carbon sources via gluconeogenesis, glyoxylate cycle, and TCA cycle (Coccetti *et al*, 2018; Hardie, 2007; Rashida *et al*, 2021). Additionally, glucose starvation activates the Rim15 kinase, which controls storage carbohydrate metabolism and stress responses (Pedruzzi *et al*, 2003; Reinders *et al*, 1998; Jaquenoud *et al*, 2012). Given the importance of regulating gluconeogenic outputs as glucose depletes, which involves extensive phosphorylation-based regulation, control by protein dephosphorylation would be anticipated for such metabolic adaptation. However, the known roles of phosphatases in carbon metabolism are limited to a few examples. The PP1 phosphatase dephosphorylates and inactivates the Snf1 kinase to enable glucose-mediated catabolite repression in glucose-replete conditions (Rubenstein *et al*, 2008; Hisamoto *et al*, 1994). Upon glucose starvation, PP2A phosphatases dephosphorylate stress-responsive transcription factors to regulate the synthesis of storage carbohydrates (Jaquenoud *et al*, 2012; Baro *et al*, 2018; Offley & Schmidt, 2019; Ariño *et al*, 2019). Given the numbers of phosphatases in any eukaryotic genome including yeast (Offley & Schmidt, 2019), it is plausible that other protein phosphatases play roles in metabolic adaptations to changing glucose environments.

Here, using a phosphatase knockout screen, we identify a role for the PP2A-like phosphatase Ppg1, in regulating metabolic adaptations in the post-diauxic phase. Using *ppg1Δ1* and Ppg1 catalytically-inactive cells, we establish the role of Ppg1 in controlling gluconeogenic flux and carbon allocations post glucose depletion. We identify that Ppg1 carries out this function via controlling the assembly of a multiprotein scaffolding complex - the Far complex. The Ppg1 and Far complex-mediated regulation of gluconeogenic outputs are critical for cells to adapt to glucose depletion in competitive growth environments where glucose gets depleted rapidly. Collectively, we identify a mechanism where cells use protein dephosphorylation to dynamically assemble a scaffolding complex on organelles as assembly hubs, through which cells can tune carbon allocations in order to adapt to fluctuating nutrient environments.

## Results

### A phosphatase deletion screen identifies Ppg1 as a regulator of storage carbohydrate metabolism

As glucose depletes, Crabtree-positive cells such as *S. cerevisiae* switch from glycolytic fermentation to gluconeogenesis and utilization of other carbon sources (Figure 1A). Our objective was to identify protein phosphatases that regulate this metabolic adaptation (Figure 1A). To address this, we generated a *S. cerevisiae* phosphatase deletion library consisting of individual deletions of 38 non-essential phosphatases (Figure S1A), and carried out a screen to identify phosphatases that regulate carbon metabolism post glucose depletion. For this screen, wild-type and phosphatase deletion mutant cells were cultured in a glucose-replete medium (YPD), and grown post glucose depletion (post-diauxic phase). For the screen output, trehalose accumulation was assessed from these mutants after 24hrs. Trehalose synthesis increases in the post-diauxic phase and is a reliable readout of a gluconeogenic state (Varahan *et al*, 2019b; Parrou, 2001) (Figure 1B). Using this approach, we identified phosphatase mutants with increased or decreased trehalose accumulation (Figure 1C, S1A). A notable ‘hit’ was the PP2A-like phosphatase Ppg1, where the loss of Ppg1 results in a significant increase in trehalose accumulation in the post-diauxic phase (Figure 1C). This increase in post-diauxic trehalose levels was considerable and suggests that Ppg1 might balance carbon metabolism upon glucose depletion. We therefore focused on deciphering role of Ppg1 in these processes. Currently, there is limited evidence for Ppg1 in regulating glycogen metabolism (Posas *et al*, 1993); and its roles or mechanisms in regulating metabolic adaptation are unknown.

**Figure 1:**
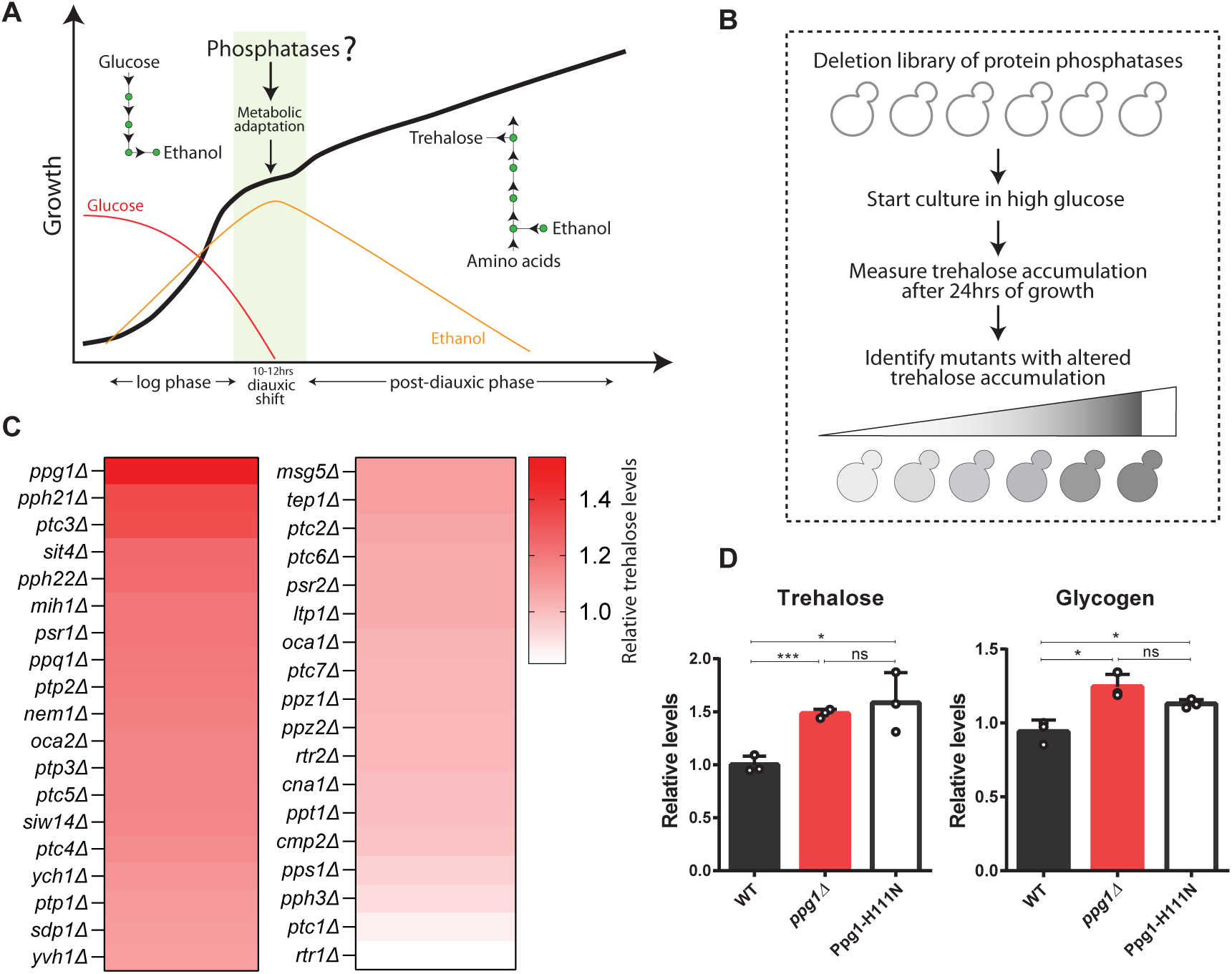
A phosphatase deletion screen identifies Ppg1 as a regulator of storage carbohydrate metabolism. A) Schematic depicting growth kinetics of *S. cerevisiae* cells in glucose-replete conditions and varying metabolic states during different phases of growth. *S cerevisiae* cells rewire their metabolism after diauxic adaptation. B) Schematic describing the experimental flow of the screen to identify protein phosphatases that regulate post-diauxic carbon metabolism. For the screen, phosphatase knockouts were grown in a glucose-replete medium, and trehalose accumulation was measured after 24hrs of growth. C) Identifying phosphatase knockouts with altered trehalose amounts. The heat map shows relative trehalose amounts in phosphatase knockouts compared to wild type cells. Trehalose accumulation was measured after 24hrs of growth in YPD medium. The mean trehalose levels was from 2 biological replicates. D) Effect of loss of phosphatase activity of Ppg1 on the accumulation of storage carbohydrates in post-diauxic phase. A catalytically inactive mutant of Ppg1-Ppg1H111N was generated. WT, *ppg1Δ1*, and Ppg1H111N cells were grown in a glucose replete medium and trehalose and glycogen were measured after 24hrs. Data represented as a mean ± SD (n=3). *P < 0.05, **P < 0.01, and ***P< 0.001; n.s., non-significant difference, calculated using unpaired Student’s t tests.

While little is known about Ppg1 functions in maintaining metabolic homeostasis, a previous study identified a role for Ppg1 in preventing mitophagy, by dephosphorylating the autophagy-regulating protein Atg32 (Furukawa *et al*, 2018b). Although the conditions in this study do not induce mitophagy, we still asked if these distinct roles were connected using Atg32 mutants. However, a loss in this mitophagy regulator had no impact on trehalose accumulation (Figure S1B), suggesting that this function of Ppg1 in controlling trehalose metabolism is independent of its role in mitophagy. We next assessed if both glycogen and trehalose (which are synthesized from a common node of glucose-6-phosphate produced by gluconeogenesis) were regulated by Ppg1. The loss of Ppg1 phosphatase showed increased glycogen accumulation in post-diauxic phase (Figure 1D), similar to the accumulation of trehalose. In order to determine if this function of Ppg1 in storage carbohydrate metabolism depends on its phosphatase activity, we generated a catalytic-dead single point mutant of Ppg1 – Ppg1H111N at the native Ppg1 chromosomal locus (Innokentev *et al*, 2020), and measured levels of trehalose and glycogen in these cells. Similar to *ppg1Δ1* cells, Ppg1H111N cells showed an increase in amounts of trehalose and glycogen in post-diauxic phase, confirming that the phosphatase activity of Ppg1 is required to control storage carbohydrate metabolism (Figure 1D). As an added control, we confirmed that this point mutation in the catalytic site did not affect steady-state levels of Ppg1 protein (Figure S1C).

Collectively these data establish that the loss of Ppg1 leads to an accumulation of storage carbohydrates in the post-diauxic phase, and this depends on Ppg1 phosphatase activity.

### Ppg1 controls gluconeogenic flux and outputs in the post-diauxic phase

Since the loss of Ppg1 increased storage carbohydrates, we asked if other gluconeogenic outputs are altered in these cells. To address this, we used quantitative, targeted LC-MS/MS to identify changes in carbon allocation to other metabolic arms. We first assessed relative amounts of these metabolites (steady-state) from WT and *ppg1Δ1* cells after 24hrs of growth in YPD. Notably, steady-state amounts of glucose-6-phosphate (G6P), fructose-6-phosphate (F6P), fructose-1,6-bisphosphate (F16BP) (which would all come from gluconeogenesis post glucose depletion) significantly increased in *ppg1Δ1* cells. The levels of UDP-Glucose – a precursor of storage carbohydrates and cell wall increased in *ppg1Δ1* cells. UDP-GlcNAc (a precursor of the chitin component of cell wall) was also higher in *ppg1Δ1* cells (Figure 2A). Upon glucose depletion, amino acids can provide carbon backbones for gluconeogenesis (Varahan *et al*, 2020). We therefore also assessed the amounts of amino acids in these cells. The steady-state amounts of multiple free amino acids significantly decreased in *ppg1Δ1* cells (Figure 2A). Collectively, these data suggest that *ppg1Δ1* cells have imbalanced carbon metabolism, where amino acids might be consumed at an increased rate fueling gluconeogenesis, and carbon flux may be directed towards synthesis of storage carbohydrates and precursors of cell wall. This was also observed in Ppg1H111N cells, confirming the role of Ppg1 phosphatase activity in regulating carbon metabolism post glucose depletion (Figure S2A).

**Figure 2:**
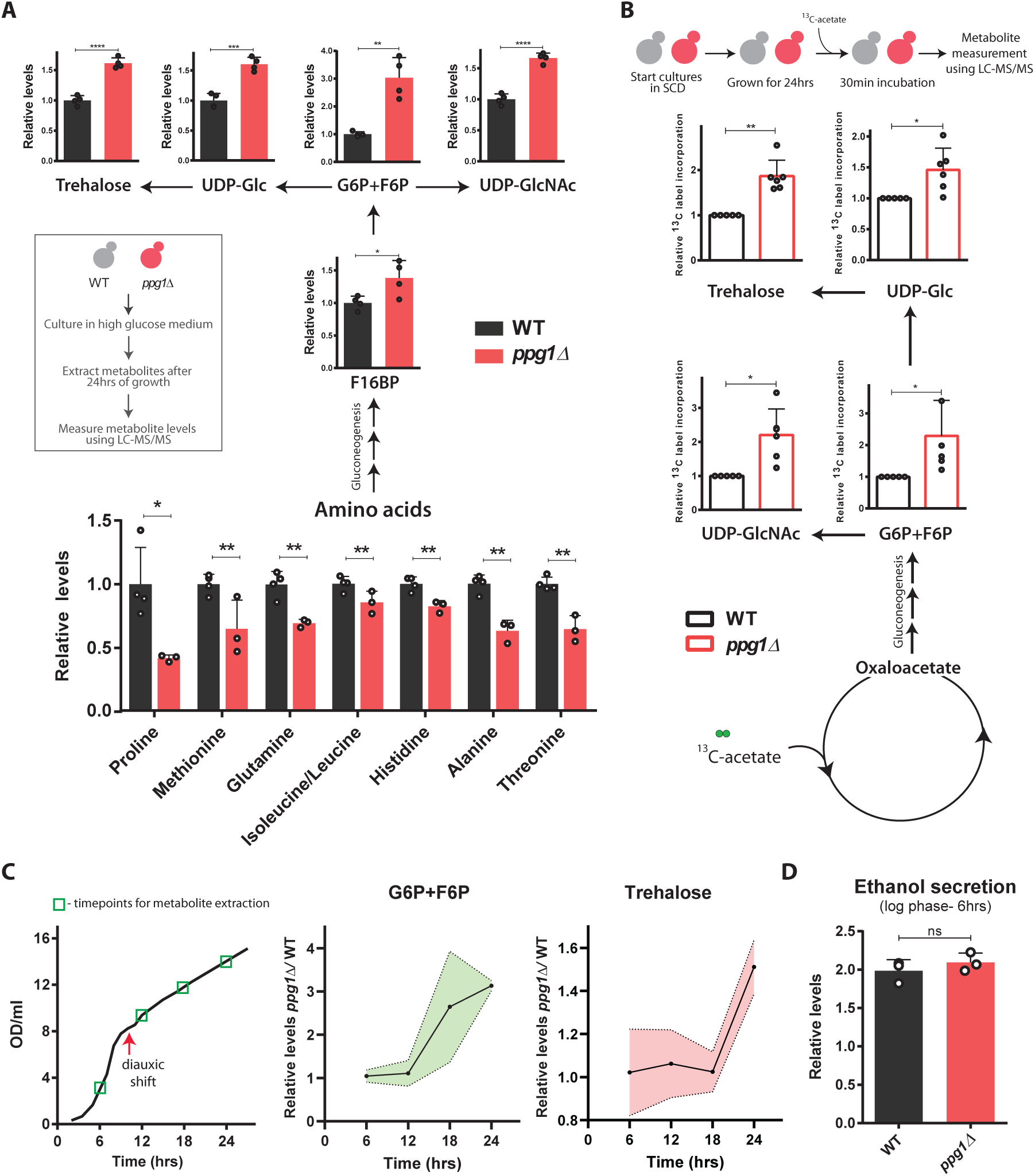
Ppg1 controls gluconeogenic outputs in the post-diauxic phase. A) Relative steady-state amounts of specific gluconeogenic intermediates, precursors of cell wall and storage carbohydrates, and amino acids in WT and *ppg1Δ1* cells after 24hrs of growth in YPD medium. Data represented as a mean ± SD (n=3). G6P, glucose-6-phosphate; F6P, fructose-6-phosphate; UDP-Glc, uridine diphosphate glucose; UDP-GlcNAc, uridine diphosphate N-acetylglucosamine. *P < 0.05, **P < 0.01, and ***P< 0.001; n.s., non-significant difference, calculated using unpaired Student’s t tests. B) Relative 13C label incorporation in gluconeogenic outputs. WT and *ppg1Δ1* cells were grown in SCD medium for 24hrs, cultures were spiked with 13C-acetate for 30mins, and 13C label incorporation in gluconeogenic outputs was measured. Data represented as a mean ± SD (n=6). *P < 0.05, **P < 0.01, and ***P< 0.001; n.s., non-significant difference, calculated using paired t tests. C) Relative steady-state amounts of specific gluconeogenic outputs from WT and *ppg1Δ1* cells during the course of growth in YPD medium. WT and *ppg1Δ1* cells were grown in YPD medium, and metabolite extraction was carried out at indicated time points. Also, see Fig. S2 D. Data represented as a mean ± SD (n=3). D) Relative ethanol production by WT and *ppg1Δ1* cells in high-glucose conditions. WT and *ppg1Δ1* cells were grown in YPD medium for 6hrs and ethanol concentration in the media was measured. Data represented as a mean ± SD (n=3). *P < 0.05, **P < 0.01, and ***P< 0.001; n.s., non-significant difference, calculated using unpaired student’s t tests.

Since steady-state metabolite levels do not always indicate changes in flux, in order to unambiguously address if carbon flux was imbalanced in *ppg1Δ1* cells, we compared carbon flux through these pathways in WT and *ppg1Δ1* cells. Specifically, we used a pulse label of 1% ^13^C2-acetate (which can directly enter gluconeogenesis) to cells in the post-diauxic phase, and estimated label incorporation into gluconeogenic intermediates, precursors of cell wall and storage carbohydrates. The relative 13C label incorporation into gluconeogenic intermediates significantly increased, as did the label incorporation into trehalose, UDP-Glc, and UDP-GlcNAc, in *ppg1Δ1* cells (Figure 2B). These data reveal a higher gluconeogenic flux, coupled with increased carbon allocation towards synthesis of storage carbohydrates and cell wall precursors in *ppg1Δ1* cells. This metabolic characterization of *ppg1Δ1* cells thus revealed a specific imbalance in carbon allocation in the post-diauxic phase.

To understand if the Ppg1-dependent metabolic regulation is specific to the post-diauxic phase, we also assessed its role in the glycolytic phase of growth in glucose-replete conditions. For this, we compared these same metabolites at different time points starting from a glucose-replete medium. In *ppg1Δ1* cells, the steady-state amounts of intermediates of glycolysis, precursors of cell wall and storage carbohydrates were unaltered compared to WT cells in the pre-diauxic phase. In *ppg1Δ1* cells, the increased accumulation of these metabolites was observed only after glucose depletion in the post-diauxic phase (Figure 2C, S2B). We also assessed the effect of Ppg1 deletion on fermentation by measuring ethanol secretion in WT and *ppg1Δ1* cells. In these conditions (prior to the diauxic shift), the amount of ethanol produced by *ppg1Δ1* cells was indistinguishable from WT cells, suggesting that loss of Ppg1 does not affect glycolytic outputs in high glucose (Figure 2D).

Finally, since the carbon allocation towards synthesis of UDP-GlcNAc increases in *ppg1Δ1* cells, we measured the levels of chitin from the cell walls of post-diauxic cells. Consistently, chitin levels increased in *ppg1Δ1* cells (Figure S2C). Increased chitin accumulation is known to sensitize cells to cell wall stress (Ram & Klis, 2006; Vannini *et al*, 1983); hence, we studied the growth of *ppg1Δ1* cells in the presence of Congo red (a cell wall stress agent). Expectedly, (and as observed earlier (Hirasaki *et al*, 2010)) the growth of *ppg1Δ1* cells was reduced in the presence of Congo red (Figure S2D).

Collectively, these results establish that Ppg1 phosphatase controls the allocation of carbon towards gluconeogenic outputs in the post-diauxic phase of growth.

### Ppg1 regulates carbon metabolism via Far complex

How might Ppg1 regulate this balance of carbon flux in glucose deplete environments? Ppg1 is a PP2A-like phosphatase. Several PP2A phosphatases themselves lack substrate specificity, and therefore function with associated proteins that facilitate the transient interaction of the phosphatase with its substrates (Slupe *et al*, 2011; Roy & Cyert, 2009). We hypothesized that identifying interacting partners of Ppg1 could help us understand its specific mechanisms of regulation. To identify proteins interacting with Ppg1, we introduced a FLAG epitope tag at C-terminus of Ppg1 at its chromosomal locus, performed a FLAG affinity purification/elution from post-diauxic cells, with wild-type (untagged) cells as control. Immunoprecipitated fractions were resolved on SDS gels and were visualized by silver staining. The protein gel revealed obvious differences in the immunoprecipitated proteins between Ppg1 and control samples (Figure 3A). The gel fragments were excised, and proteins were identified using mass spectrometry. A total of 33 and 24 proteins were identified in Ppg1 immunoprecipitation fractions (replicate 1 and 2 respectively). The list of proteins identified was filtered manually by eliminating contaminant proteins and proteins found in control fractions (non-specific interactors), and retaining only those present in both biological replicates. The proteins uniquely identified in the Ppg1 immunoprecipitation fraction as Ppg1 interactors included Tpd3, Far11, and Ssa1 (Figure 3A). Tpd3 is a regulatory sub-unit of the PP2A phosphatase and is therefore expected, while Ssa1 is a chaperone of Hsp70 family. Hence, these two interactions were not further considered.

**Figure 3:**
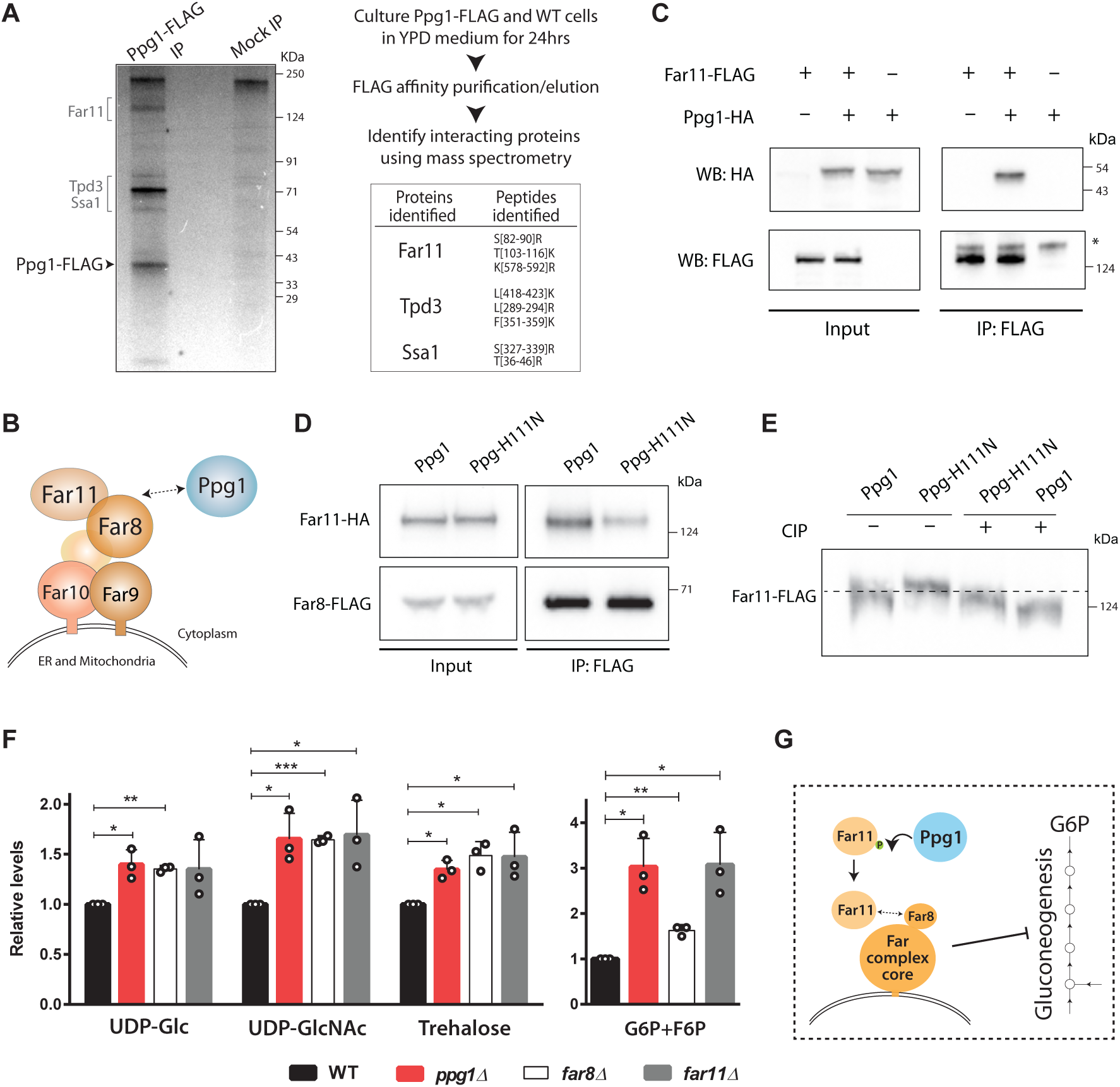
Ppg1 regulates gluconeogenic carbon allocations via the Far complex. A) Identification of interacting partners of Ppg1. Lysates of WT and Ppg1-FLAG cells were subjected to FLAG affinity purification, immunopurified fractions were separated on SDS-PAGE followed by silver staining, and the immunopurified fractions were analyzed by mass spectrometry. The peptides identified for the proteins specifically interacting with Ppg1 are listed. The experiments were performed using 2 biological replicates. B) Schematic describing the composition of Far complex. C) Ppg1 interacts with Far11. FLAG-tagged Far11 was immunoprecipitated from cells after 24hrs of growth in YPD medium and co-immunopurified HA-tagged Ppg1 was detected using western blotting. * denotes a non-specific binding of the antibody. A representative image is shown (n=2). D) Requirement of Ppg1 activity for the interaction between Far11 and Far8. WT and Ppg1H111N cells containing HA-tagged Far11 and FLAG-tagged Far8 were cultured in YPD medium for 24hrs. Far8-FLAG was immunoprecipitated from these cells, and co-immunopurified Far11-HA was detected. A representative image is shown (n=2). E) Regulation of Far11 post-translational modifications by Ppg1. WT and Ppg1H111N cells containing endogenously tagged Far11 with 3xFLAG epitope were cultured in YPD medium for 24hrs. Protein precipitation was carried out and Far11 mobility was monitored on a 7% SDS-PAGE gel. For the phosphatase treatment, protein precipitates were dissolved in lysis buffer and treated with calf-intestinal phosphatase (CIP). A representative image is shown (n=3). F) Relative steady-state amounts of specific gluconeogenic outputs from WT, *ppg1Δ1*, *far8Δ1*, and *far11Δ1* cells after 24hrs of growth in YPD medium. Data represented as a mean ± SD (n=3). *P < 0.05, **P < 0.01, and ***P< 0.001; n.s., non-significant difference, calculated using paired t tests. G) Schematic describing Ppg1 regulating Far complex assembly to control gluconeogenic carbon flux.

The identification of Far11 protein in this context of metabolic adaptation was however unexpected, since there is no reported role for this protein in metabolic homeostasis. Far11 is a component of the Far complex - a multiprotein scaffolding complex consisting of six subunits (Far11, Far8, Far3, Far7, Far9, and Far10 subunits) (Kemp & Sprague, 2003) (Figure 3B), where the interaction between Far8 and Far11 completes the complex assembly (Pracheil & Liu, 2013). Currently, this complex is known to prevent mitophagy during nitrogen starvation, and also functions in pheromone-mediated cell cycle arrest and TORC2 signaling (Pracheil *et al*, 2012). Previous studies have found that Ppg1 phosphatase can interact with the Far complex (Furukawa *et al*., 2018). However, none of these roles are related to the homeostatic regulation of carbon metabolism.

Therefore, to first better understand interactions or relationships between Far complex and Ppg1, we first biochemically characterized the interaction between the Far complex and Ppg1 phosphatase using co-immunoprecipitation. Specifically, we immunopurified Far11 from cells growing in post-diauxic phase and assessed the association of Ppg1 with immunopurified Far11 (Figure 3C). The immunopurified Far11 specifically co-precipitated Ppg1 in this experiment (Figure 3C). We next asked if the phosphatase catalytic activity of Ppg1 was required for Far complex assembly, and performed similar co-immunopurification experiments using WT and Ppg1H111N cells. Consistently, we observed that the co-immunoprecipitation of Far11 with Far8 is notably reduced in Ppg1H111N cells (Figure 3D). This suggests that the interaction between Far11 and Far8 depends on the phosphatase activity of Ppg1 (Figure 3D). These data are consistent with a previous report suggesting that Ppg1 is required for Far complex assembly (Innokentev *et al*, 2020).

Since Ppg1 activity was critical for interaction between components of the Far complex, we asked if Far11 or Far8 were phosphorylated, in a Ppg1-dependent manner. To assess this, we investigated changes in the electrophoretic mobility of Far8 and Far11 on SDS-PAGE gels. Changes in the electrophoretic mobility of a protein are a well-established read-out of protein phosphorylation/dephosphorylation (Lee *et al*, 2013). We compared the electrophoretic mobility of Far8 and Far11 in WT and Ppg1H111N on SDS-PAGE gels. While the mobility of Far8 was not altered in Ppg1H111N cells, Far11 mobility in Ppg1H111N cells was reduced compared to WT (Figure 3E, S3A). This would be consistent with Ppg1-dependent dephosphorylation of Far11, and therefore increased phosphorylation of Far11 in Ppg1H111N cells. To further examine this possibility, we treated protein extracts with alkaline phosphatase and monitored Far11 electrophoretic mobility. The phosphatase treatment of protein extracts from post diauxic wild-type cells resulted in reduced Far11 mobility (Figure 3E). This suggests that Far11 is phosphorylated in post-diauxic WT cells (Figure 3E). Additionally, the phosphatase-treated Far11 from Ppg1H111N cells show further reduced electrophoretic mobility compared to phosphatase-treated Far11 from WT cells. Collectively, these data suggest that Far11 is post-translationally modified, and this modification depends on Ppg1 phosphatase activity. These data indicate that Ppg1 controls the assembly of the Far complex, via the post-translational regulation of Far11.

Therefore, we asked if this (Ppg1-dependent assembly of) Far complex was required to regulate gluconeogenic outputs. We compared levels of gluconeogenic intermediates, precursors of storage carbohydrates and cell wall in post-diauxic phase from *far8Δ1* and *far11Δ1* cells. Similar to *ppg1Δ1* cells, the amounts of all these metabolites increased in *far8Δ1* and *far11Δ1* cells (Figure 3F, S3B). Furthermore, the deletion mutants of components of Far complex were sensitive towards Congo red (Figure S3C), phenocopying the *ppg1Δ1* cells. Collectively, these data find that the loss of the Far complex phenocopies the loss of Ppg1, and the assembled Far complex is required to appropriately apportion carbon allocations in post-diauxic cells.

Summarizing, these data show that the Ppg1 phosphatase activity regulates the assembly of Far complex, which thereby regulates carbon flux towards gluconeogenic outputs important in post-diauxic cells (Figure 3G).

### Far complex tethering and not specific localization is required for gluconeogenic regulation

Interestingly, the Far complex is present in two sub-cellular localizations within a cell. One subpopulation localizes to the outer membrane of ER and other to the mitochondrial outer membrane (Innokentev *et al*, 2020; Onishi *et al*, 2023), both functioning as cytosol-facing complexes. We wondered if any one of the subpopulations of the Far complex was required for this regulation of carbon flux (Figure 4A). Therefore, we decided to study the role of each subpopulation of the Far complex in the context of this function.

**Figure 4:**
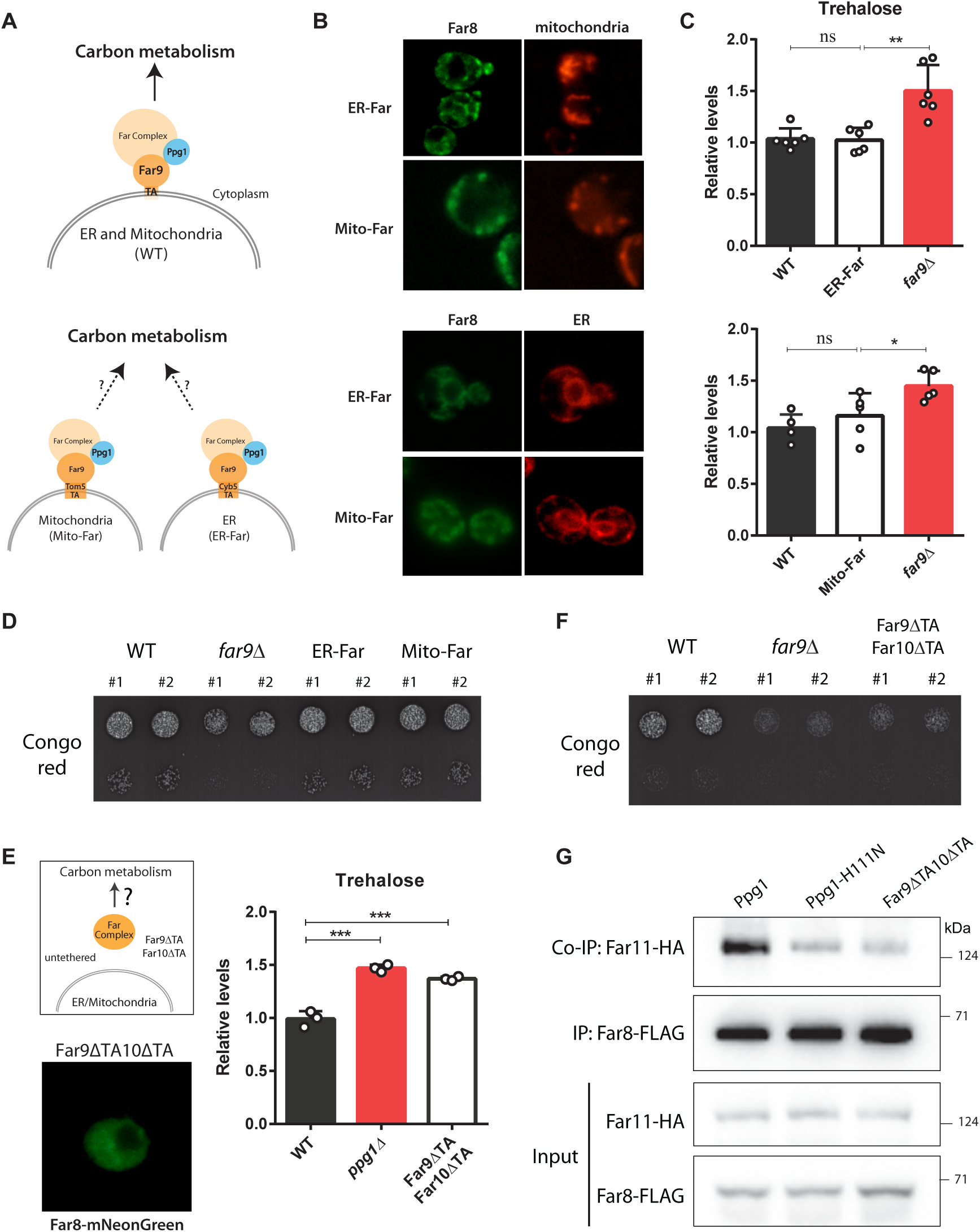
Far complex tethering and not specific subcellular localization enables appropriate carbon allocations. A) Schematic describing two sub-populations of Far complex and the possibility of a specific subpopulation involved in regulating carbon metabolism. The two sub populations of Far complex are present at mitochondrial and ER outer membranes. The strains Mito-Far and ER-Far were constructed, and in these strains Far complex localizes specifically to mitochondria and ER respectively. B) The Mito-Far and ER-Far strains show distinct mitochondrial and ER localization of the Far complex. The Mito-Far and ER-Far cells were grown in YPD medium for 24hrs and analyzed by fluorescent microscopy. Far8-mNeonGreen was used to visualize Far complex. Sec63-mCherry was used to visualize ER. Mitochondria were visualized using mitotracker red CMXRos. Representative images are shown. C) Effect of targeting of Far complex to ER and mitochondria on trehalose accumulation. WT, Mito-Far, ER-Far, and *far9Δ1* cells were cultured in YPD medium for 24hrs and trehalose accumulation was measured from these cells. Data represented as a mean ± SD (n=6 for ER-Far and n=5 for Mito-Far). *P < 0.05, **P < 0.01, and ***P< 0.001; n.s., non-significant difference, calculated using unpaired t tests. D) The growth of Mito-Far and ER-Far cells in presence of Congo red. A serial dilution growth assay was carried out in presence of Congo red using WT, Mito-Far, ER-Far, and *far9Δ1* cells. Congo red was used at a final concentration of 400 μg/ml. The images were taken after 48hrs of growth. A representative image is shown (n=2). E) Role of membrane tethering of Far complex on trehalose accumulation. The TA domains of Far9 and Far10 were deleted to disrupt the membrane tethering of Far complex (Far9*⊗*1TA10*⊗*1TA cells). WT, *far9Δ1*, and Far9*⊗*1TA10*⊗*1TA cells were cultured in YPD medium for 24hrs and trehalose accumulation was measured from these cells. Data represented as a mean ± SD (n=3). *P < 0.05, **P < 0.01, and ***P< 0.001; n.s., non-significant difference, calculated using unpaired t tests. F) The growth of Far9*⊗*1TA10*⊗*1TA cells is attenuated in the presence of Congo red. A serial dilution growth assay was carried out in the presence of Congo red using WT, *far9Δ1*, and Far9*⊗*1TA10*⊗*1TA cells. Congo red was used at a final concentration of 400 μg/ml. The images were taken after 48hrs of growth. A representative image is shown (n=2). G) The interaction between Far8 and Far11 in Far9*⊗*1TA10*⊗*1TA cells. WT, Ppg1H111N, and Far9*⊗*1TA10*⊗*1TA cells expressing HA-tagged Far11 and FLAG-tagged Far8 were cultured in YPD medium for 24hrs. Far8-FLAG was immunoprecipitated from these cells, and co-immunoprecipitated Far11-HA was detected. A representative image is shown (n=3).

The Far9 and Far10 proteins contain tail-anchor (TA) domains required to tether the complex to the ER or mitochondrial outer membrane. To determine which of the ER or the mitochondria-localized Far regulates post-diauxic carbon flux, we genome-engineered two strains with distinct Far localizations to either the ER or mitochondrial outer membrane. For these strains, we replaced the native TA domain of Far9 with the TA domains of Tom5 or Cyb5, which will localize the Far complex specifically to only the mitochondrial surface or only the ER surface, respectively. As anticipated, Far8-mNeonGreen from Far9-Cyb5 (ER-Far) and Far9-Tom5 (Mito-Far) cells showed exclusive ER or mitochondrial localization, respectively (Figure 4B). As readouts of wild-type Far complex function, we measured trehalose accumulation in post-diauxic phase from these mutant cells. Surprisingly, both the Far9-Cyb5 (ER-Far) as well as the Far9-Tom5 (Mito-Far) strains retained trehalose accumulation, similar to WT cells (Figure 4C), and unlike the *far9Δ1* or *ppg1Δ1* cells. This suggested that carbon allocations remained unaltered in these cells with mitochondrial or ER surface localized Far complex (Figure 4C). Furthermore, these cells were not sensitive towards Congo red (Figure 4D, S4A), indicating that the cell wall composition of Far9-Cyb5 and Far9-Tom5 cells remained unaltered. These data therefore suggest that the specific localization of the Far complex was not critical to control this function of regulating carbon metabolism.

Therefore, we hypothesized that perhaps the Far complex required membrane tethering on a suitable cytosol-facing surface for its complete assembly. In such a scenario, the tethering of the complex on an available cytosol-facing membrane surface, along with the regulated dephosphorylation by Ppg1, would determine the complete Far complex assembly, and thereby regulate carbon metabolism. To disrupt the membrane tethering of Far complex, we first removed the TA domain of only Far9. Surprisingly, these Far9*⊗*1TA cells retained a membrane-localized Far8 protein (Figure S4B) and showed similar trehalose accumulation as WT cells (Figure S4C). Therefore, the deletion of the TA domain of Far9 alone is insufficient to disrupt the membrane tethering of the Far complex. We next removed the TA domains of both Far9 and Far10 within a single strain. The cells with this TA domain deletion in both Far9 and Far10 (Far9*⊗*1TA10*⊗*1TA) now showed cytosolic localization of Far8-mNeonGreen, and the membrane localization of Far complex was completely abolished (Figure 4E). We assessed trehalose accumulation and sensitivity to Congo red in these strains. Notably, the Far9*⊗*1TA10*⊗*1TA cells accumulated trehalose and glycogen in the post-diauxic phase, (Figure 4E, S4D), and were sensitive to Congo red (Figure 4F, S4E), similar to *ppg1Δ1* cells. Collectively, these data reveal that the loss of tethering to both the ER and mitochondrial membrane disrupted the Far complex function and that the membrane tethering of the Far complex is required for appropriate carbon flux regulation in post-diauxic cells.

The simplest explanation for this reduced Far complex function in Far9*⊗*1TA10*⊗*1TA cells is that the tethering of Far to an available cytosol-facing membrane surface is required for the complete assembly of the complex. To therefore test if the Far complex assembly is disrupted after the loss of membrane tethering, we examined the interaction between Far11 and Far8 using co-immunoprecipitation. In Far9*⊗*1TA10*⊗*1TA cells, the interaction between Far11 and Far8 was greatly reduced compared to the wild-type cells (Figure 4G), indicating that membrane tethering of Far complex is required for complete complex assembly.

These data collectively show the requirement of membrane tethering for the efficient assembly of the Far complex, and this assembled complex regulates carbon allocations towards gluconeogenic outputs. The specific subcellular localization of the Far complex to either the ER or mitochondria is itself not important for controlling carbon allocation.

### Glucose availability controls levels of the Far complex

Since the Ppg1-Far complex mediated regulation of carbon metabolism is specific to the post diauxic phase, how responsive were the Far complex or Ppg1 itself to glucose levels? For this, we first measured amounts of Ppg1 and the components of Far complex during the course of growth, in pre-diauxic and post-diauxic phase cells. The amounts of Ppg1 remained constant during the course of growth in YPD medium (Figure 5A). In contrast, the levels of Far11 increased after 24hrs of growth in YPD medium (Figure 5B). From these findings, it can be inferred that the amount of Far complex itself increases in post-diauxic phase. We further examined the interaction between Far8 and Far11 from cells in the pre-diauxic and post diauxic phases. In both pre-diauxic phase and post-diauxic phase cells, the interaction between Far11 and Far8 was retained (Figure S5A). However, using the electrophoretic mobility-based assay shown earlier, we observed that Far11 shows an electrophoretic mobility shift after glucose depletion (after 12 and 24hrs of growth in YPD) (Figure 5C). This data suggests that Far11 gets post-translationally modified as glucose depletes. The increased amounts and post-translational regulation of Far11 were also observed in Ppg1-H111N cells (Figure 5C), indicating an alternative Ppg1-independent regulation of the Far complex in response to glucose depletion.

**Figure 5:**
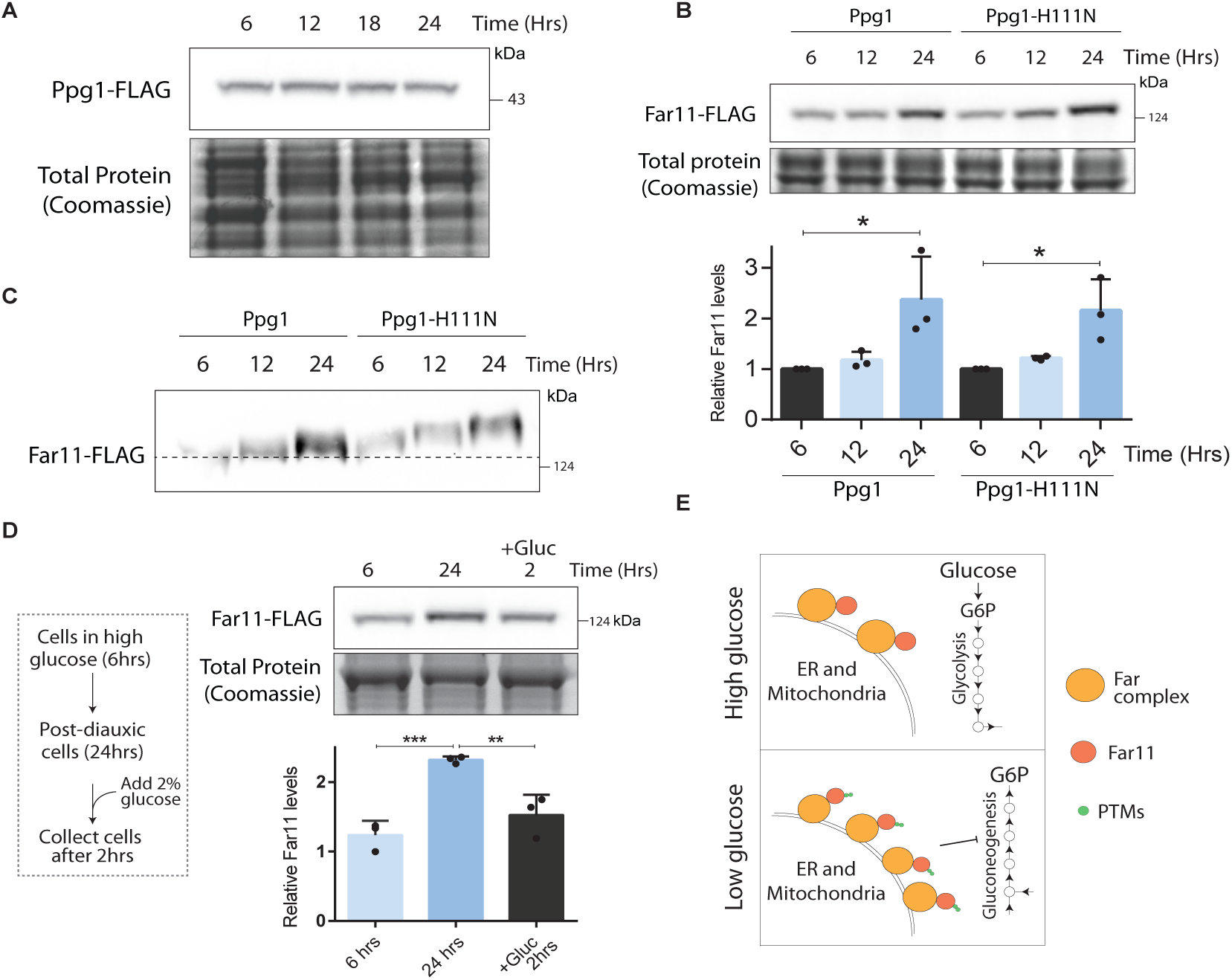
Glucose availability controls amounts of the Far complex. A) Levels of Ppg1 protein during the course of growth. Cells with Ppg1 tagged with FLAG epitope at the C terminal were cultured in YPD medium. Cells were collected at indicated time points and levels of Ppg1-FLAG were measured by western blotting. A portion of the gel was Coomassie stained and used as a loading control. A representative image is shown (n=3). B) Levels of Far11 protein during the course of growth. The WT and Ppg1H111N cells containing endogenously tagged Far11 with 3xFLAG epitope were cultured in YPD medium. Cells were collected at indicated time points and levels of Far11-FLAG were measured by western blotting. A portion of the gel was Coomassie stained and used as a loading control. Western band quantification was done using ImageJ software. A representative image is shown (n=3). *P < 0.05, **P < 0.01, and ***P< 0.001; n.s., non-significant difference, calculated using paired t tests. C) Far11 post-translational modifications during the course of growth. WT and Ppg1H111N cells containing endogenously tagged Far11 with 3xFLAG epitope were cultured in YPD medium. Cells were collected at indicated time points and Far11 mobility was monitored on a 7% SDS-PAGE gel. A representative image is shown (n=3). D) Effect of glucose availability on amounts of Far11. Cells expressing FLAG-tagged Far11 were cultured in YPD medium. After 24hrs of growth cells were diluted in the spent medium and 2% glucose was added. Cells were collected after 2hrs and levels of Far11 were measured by western blotting. A portion of the gel was Coomassie stained and used as a loading control. Western band quantification was done using ImageJ software. A representative image is shown (n=3). *P < 0.05, **P < 0.01, and ***P< 0.001; n.s., non-significant difference, calculated using unpaired t tests. E) Schematic describing the effect of glucose availability on Far complex amounts.

Finally, if glucose availability regulated the amounts of the Far complex, one possibility would be that the addition of glucose to post-diauxic cells would reduce Far complex amounts. To test this, we grew cells in high glucose and once cells reached post-diauxic phase, we added glucose and compared the levels of Far11 protein. Glucose addition to post-diauxic cells reduced the amounts of Far11 to that observed in pre-diauxic phase of growth (Figure 5D). Furthermore, the amounts of Far8 also were reduced after addition of glucose to post-diauxic cells (Figure S5C). Together, we infer that the activity and amounts of Ppg1 are constitutive, but the amounts of the Far proteins are glucose-responsive (Figure 5E). The increased amounts of this assembled Far complex therefore modulate gluconeogenic outputs in post diauxic cells.

### Ppg1-Far complex mediated carbon flux regulation enables cells to adapt to glucose depletion

Having identified this role of the Ppg1 phosphatase and Far complex in modulating post diauxic carbon allocations towards gluconeogenesis, we asked if this Ppg1-Far mediated regulation enables cells to better adapt to glucose depletion. To address this, we designed a competitive growth-fitness experiment using WT and *ppg1Δ1* cells, as illustrated in Figure 6A. For this, we used WT and *ppg1Δ1* cells constitutively expressing mCherry and mNeonGreen fluorophores respectively to quantify different cells (see methods for details). To specifically address if Ppg1 enabled competitive growth in glucose-depleting environments, fluorescently labelled WT and *ppg1Δ1* cells were started in equal proportions and grown in a glucose-replete medium for 24hrs (i.e., post-diauxic phase) and then shifted to a glucose-replete medium, and this process of transfers was repeated daily, and the relative proportion of WT and *ppg1Δ1* cells were estimated (design in Figure 6A). Notably, the fraction of WT cells substantially increased with each transfer (Figure 6B), suggesting that Ppg1-dependent regulation enables cells to adapt to glucose-depleting environments. In control experiments, competitive growth was compared in glucose-replete environments, where WT and *ppg1Δ1* cells were grown in a glucose-replete medium, and cells were transferred to a fresh glucose-replete medium continuously every 6 hours before cells reached the diauxic phase (design in Figure 6A). Contrastingly, in this context, the relative proportion of WT and *ppg1Δ1* cells remained similar after transfers, indicating that Ppg1 function did not affect growth fitness in high glucose (Figure 6C). Finally, we further examined the effect of loss of Ppg1 on steady-state batch culture growth, starting from a glucose-replete medium. The loss of Ppg1 did not affect growth in the glucose-replete log phase, but after cells entered the post-diauxic (glucose-depleted) phase, *ppg1Δ1* cells showed reduced growth, and a reduction in biomass accumulation (Figure S6A). Collectively, these data reveal that Ppg1 enables adaptation and competitive growth-fitness after glucose depletion.

**Figure 6:**
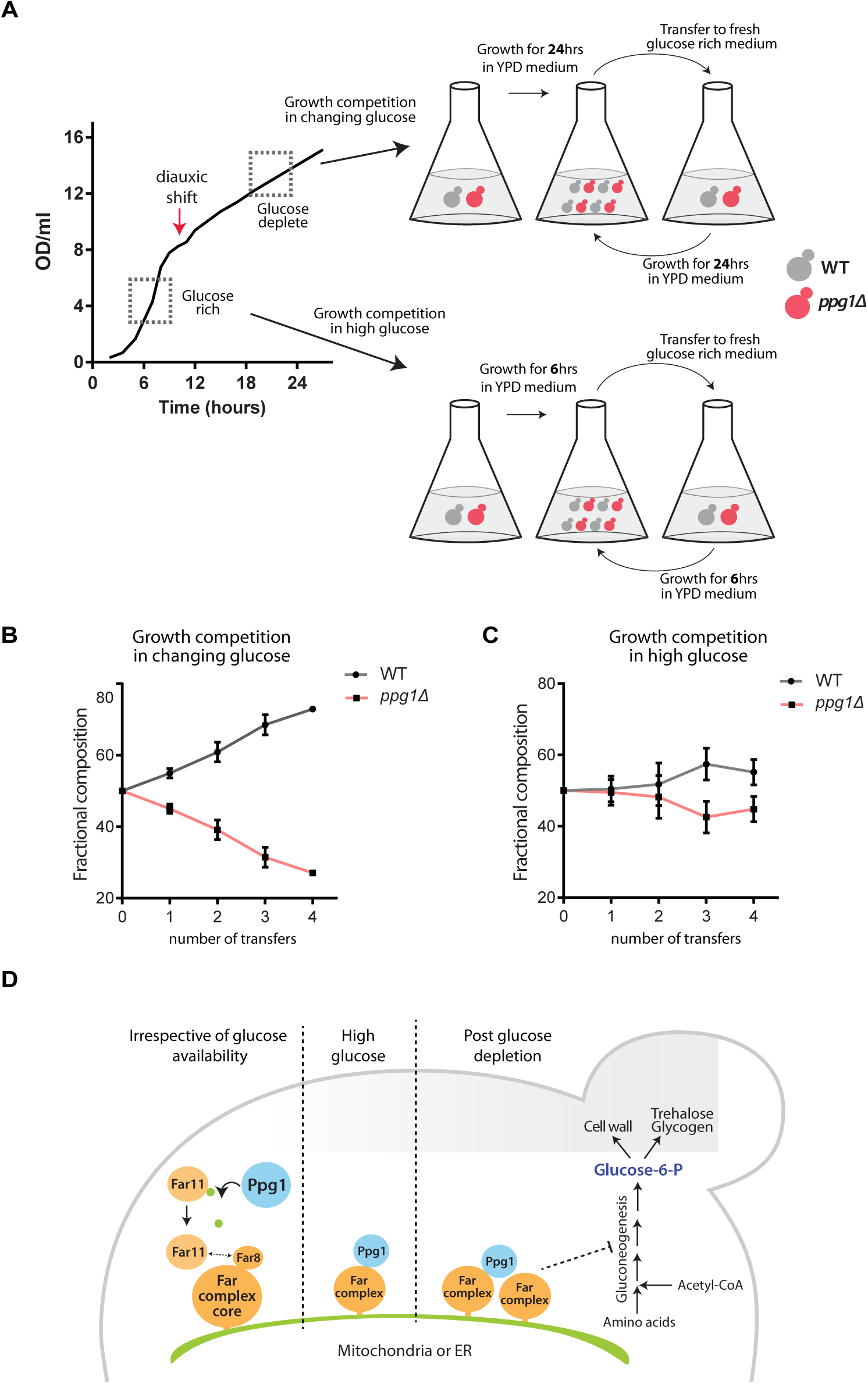
Ppg1-Far complex mediated carbon flux regulation enables cells to adapt to glucose depletion. A) Schematic describing the experimental details of growth competition experiment. WT cells expressing mCherry and *ppg1Δ1* cells expressing mNeonGreen were mixed in equal proportion. These cells were grown in YPD medium and after growth for the indicated time were shifted to fresh YPD medium. The number of WT and *ppg1Δ1* cells were calculated by fluorescent microscopy. B) Growth competition between WT and *ppg1Δ1* cells in changing glucose conditions. Data represented as a mean ± SD (n=3). C) Growth competition between WT and *ppg1Δ1* cells in glucose-replete conditions. Data represented as a mean ± SD (n=3). D) Proposed model. Ppg1 phosphatase controls the assembly of the Far complex irrespective of glucose availability. Glucose availability regulates amounts of Far complex. Ppg1-Far complex controls carbon flux and gluconeogenic outputs specifically in the low glucose.

Summarizing, we find that Ppg1 controls the assembly of the Far complex, which maintains the appropriate allocation of carbon towards gluconeogenic, post-diauxic metabolism. This Ppg1-Far mediated regulation of carbon allocation allows cells to adapt and grow competitively post glucose depletion (Figure 6C).

## Discussion

In this study, we discover a role for the PP2A-like phosphatase Ppg1 in modulating carbon allocations, which enables cells to adapt and sustain competitive growth upon glucose depletion (Figure 6D). Ppg1 inhibits the gluconeogenic carbon flux and allocation of carbon to gluconeogenic outputs as glucose depletes and cells enter the post-diauxic phase. Ppg1 mediates this function through an unexpected mechanism, where it controls the assembly of a large scaffolding protein complex, the Far complex (Figure 6D). The assembly of the Far complex requires the Ppg1 phosphatase activity, and the Ppg1-mediated Far complex assembly determines appropriate gluconeogenic outputs (Figure 6D).

While the assembly of the Far complex is itself regulated by the phosphatase activity, the Far complex amounts also increase upon depleting glucose. This exemplifies the use of reversible phosphorylation to create a dynamic scaffolding complex assembly in order to modulate metabolic states. While scaffolding proteins can integrate various biological processes, the spectrum of regulatory possibilities they enable have not been fully explored (DiRusso *et al*, 2022). Scaffolding assemblies can act as molecular facilitators to enable signaling proteins to transiently interact with correct partners, and also reduce non-specific interactions. For example, the presence of a scaffold in MAPK signaling increases the effective concentration of signaling components by nearly 3000-fold (Zeke *et al*, 2009). A scaffold could therefore modulate signal amplitude or intensity, both of which could be critical in metabolic adaptations. Such a mechanism allows cells to contextually tune outputs, without altering the signaling regulator (for example, the phosphatase) itself. A recent kinase-dependent example of such a scenario is where the yeast AMP-activated kinase (Snf1) phosphorylates Kog1, the scaffolding protein of TOR complex, leading to the disassembly of the TOR complex in extreme glucose starvation (Hughes Hallett *et al*, 2015; Prouteau *et al*, 2017; Sullivan *et al*, 2019). A parallel example is the contextual modulation of the activity of Snf1 in a TORC1 kinase independent manner, to regulate carbon allocations as glucose depletes (Rashida *et al*, 2021). In this study, the Ppg1 phosphatase functions to assemble a scaffolding complex, which would create localized pools of signaling that would regulate metabolic outputs, likely by facilitating context-dependent, specific interactions with substrates. All these examples rely on the formation of localized signaling complexes, which themselves do not have enzymatic activity, but alter metabolic outputs. We can therefore imagine scenarios in environments characterized by fluctuations in the nutrients, where dynamic molecular assemblies can contextually tune signaling and metabolic outputs, to enable competitive growth.

In addition to this identified role for the Far complex in metabolic adaptation, it also participates in other processes such as TORC2 signaling, pheromone response, and mitophagy (Furukawa *et al*, 2018b; Pracheil *et al*, 2012; Kemp & Sprague, 2003). It is unclear how the Far complex might enable signal-specificity, and distinguish between different outputs, based on distinct inputs. Our study opens the possibility that the organelle-specific localization and post-translational regulation of Far complex could enable contextual regulation of outputs. Here, while the Far11 gets modified as glucose depletes and cells enter the post-diauxic phase, the proteome itself is distinct in composition as compared to a glucose-replete environment. The modifications that decorate Far11 specifically in post-diauxic cells might enable it to interact transiently with proteins that regulate post-diauxic metabolism, or assemble signaling hubs where a phosphatase or kinase could encounter a specific substrate. In order to understand how dynamically assembled scaffolds with varying localizations and modifications can regulate homeostatic outputs such as metabolic adaptations, we require new chemical biology approaches that stabilize low-affinity protein-protein interactions, or substrate-trapping mutants to identify transient substrates that are brought together by such signaling hubs (Qin *et al*, 2021). This remains a key challenge in the context of protein phosphatases, which naturally interact with substrates with low affinities (Bonham *et al*, 2023).

Finally, homologs of the Far complex have been found in diverse eukaryotes and are known as striatin-interacting phosphatase and kinase (STRIPAK) complexes (Kück *et al*, 2016). The STRIPAK complexes regulate Hippo signaling, MAPK signaling, embryonic development, and so on (Kück *et al*, 2019; Nordzieke *et al*, 2015; Elramli *et al*, 2019). Many STRIPAK components and Far complex components are evolutionarily conserved (Kück *et al*, 2016). However, the kinases or phosphatases that associate with or regulate STRIPAK complexes remain largely unknown, and it remains unclear as to how these complexes function as signaling hubs. Therefore, identifying regulators of the Far or STRIPAK complexes, and their downstream effectors are exciting areas of future inquiry. Through the Far complex functions, it would be possible to uncover general mechanisms of how signaling hubs can assemble on available surfaces inside the cell, such as the ER or mitochondrial membrane surface. Multiple lines of inquiry now reveal that signaling pathways such as the TORC1 pathway have contextual outputs from localized signaling hubs (Hatakeyama & De Virgilio, 2019; Chen *et al*, 2021). Such localized signaling hubs have not been explored in the context of homeostatic regulation of metabolic outputs. Our work identifying the Ppg1-mediated Far complex assembly exemplifies how a localized signaling hub could modulate metabolic outputs. This advances our basic understanding of how cells tune metabolism using localized signaling hubs, in order to adapt to changing nutrient environments. Such understanding will also allow us to optimize our use of cells as factories for metabolic engineering.

## Methods

### Yeast strains, media and growth conditions

The prototrophic CEN.PK, haploid “a” strain of *S.cerevisiae* was used in all the experiments. The strains with gene deletions and chromosomally tagged proteins were generated as described in (Longtine *et al*, 1998). For all the experiments, cells were cultured in YPD medium (1% yeast extract, 2% peptone, and 2% glucose) unless otherwise mentioned. For solid agar plates, YPD media was supplemented with 2% granular agar. For all the experiments, cells were grown overnight in YPD medium, and this primary culture was used to start a secondary culture at OD600 of ∼0.2. All the cultures were incubated at 30°C/240 rpm. For the metabolic flux experiment, synthetic medium supplemented with amino acids (SCD - yeast nitrogen base without amino acids, all amino acids 2mM each with 2% glucose) was used.

### Serial dilution spot growth assay

For spot assay, ∼1 OD600 cells of respective strains were collected and washed with water. Serial dilutions were made in water and 5 μl of cell suspension was spotted on respective agar plates; growth was monitored for 48hrs. Congo red was used at a final concentration of 400 μg/ml.

### CRISPR based mutagenesis

Cells were first transformed with a plasmid expressing Cas9 (Addgene plasmid 43802) (Dicarlo *et al*, 2013). Guide RNAs (gRNAs) targeting genomic loci to be mutated were cloned in a plasmid with gRNA scaffold. The Cas9-expressing cells were transformed with gRNA plasmid and homology repair (HR) fragments. The cells were plated on YPD plates with selection drugs and the correct clones were confirmed using Sanger sequencing. The HR fragments used for transformation were synthesized.

### Western blot analysis

The cells were collected and protein extraction was carried out as described in (Rashida *et al*, 2021). Briefly, approximately ∼10 OD600 cells were collected by centrifugation and total protein was precipitated and extracted using 10% trichloroacetic acid and resuspended in SDS-glycerol buffer. The extracted proteins were estimated by BCA (bicinchoninic acid) protein assay kit. The samples were normalized for protein amounts and an equal amount of protein from all the samples were run on 4-12% bis-tris gels (Invitrogen, NP0336BOX). For the Far11 mobility shift assay, 7% SDS-PAGE gel was used. The relevant portion of gel was cut and used for transfer and blotting. The lower portion was stained with Coomassie blue for protein loading normalization. The blots were developed using the following antibodies: anti-FLAG raised in mouse (1:2000; Sigma-Aldrich, F1804), anti-HA raised in mouse (1:2000; Sigma-Aldrich, 11583816001), anti-HA raised in rabbit (1:2000; Sigma-Aldrich, H6908), anti-mouse horseradish peroxidase (HRP)-conjugated antibody (1:4000; Cell Signaling Technology, 7076S), anti-rabbit HRP-conjugated antibody (1:4000; Cell Signaling Technology, 7074S). For chemiluminescence detection, western bright ECL HRP substrate (Advansta, K12045) was used. Alkaline phosphatase treatment was performed as described in (Onishi *et al*, 2023; Rashida *et al*, 2021).

### Immunoprecipitation and co-Immunoprecipitation

Immunoprecipitation was carried out as described in (Rashida *et al*, 2021). Cells (∼50 OD600) were collected by centrifugation, the pellet was flash-frozen in liquid nitrogen and stored at - 80°C. The pellet was resuspended in lysis buffer (50 mM HEPES buffer (pH 7.0), 50 mM NaF, 10% glycerol, 150 mM KCl, 1 mM EDTA, 2 mM sodium orthovanadate, 2 mM phenylmethylsulfonyl fluoride, 0.1 mM leupeptin, 2 mM pepstatin, and 0.25% Tween 20) and cell lysis was carried out by bead-beating. The supernatant was collected by centrifugation and was precleared by incubating with Dynabeads protein G beads (Invitrogen, 10004D). The beads were pulled down using DynaMag (Invitrogen, 12321D) and the precleared lysates were incubated with Dynabeads protein G beads conjugated with anti-FLAG antibody. The suspension was incubated for 2hrs at 4°C with gentle rotation. The beads were pulled down using DynaMag and the immunoprecipitated proteins were eluted by incubating with FLAG peptide for 40 minutes. The immunoprecipitated proteins were detected by Western blotting using appropriate antibodies.

### Mass spectrometry analysis

The cells expressing Ppg1 tagged with 3xFLAG epitope and untagged cells were cultured in YPD medium for 24hrs (post-diauxic phase). Cells were collected and lysed as mentioned above. The immunopurified fraction was run on SDS-PAGE gel, the gel pieces were cut and subjected to in-gel digestion. Specifically, the gel pieces were incubated with dithiothreitol followed by incubation with iodoacetamide. The alkylated proteins were digested using Trypsin (G biosciences) for 12hrs at 37°C. Trypsin-digested peptides were dissolved in 0.1% formic acid and were analyzed on Thermo EASY-nLC™ 1200nano System coupled to a Thermo Scientific Orbitrap Fusion Tribrid Mass Spectrometer. The experiment was performed with 2 biological replicates. Data analysis was performed using Mascot distiller. Searches were conducted using 10 ppm peptide mass tolerance, product ion tolerance of 0.6 Da resulting in a 1% false discovery rate. Mascot score and emPAI values were used to compare protein abundance (Ishihama *et al*, 2005).

### Trehalose and Glycogen measurements

Trehalose and glycogen measurements were carried out as described in (Gupta *et al*, 2019; Rashida *et al*, 2021). Briefly, 10 OD600 cells were collected and lysed by incubating with 0.25M Na2CO3 at 98°C for 4hrs. The pH of solution was adjusted to 5.2 by the addition of appropriate amounts of 1M acetic acid and 0.2M CH3COONa. For trehalose estimation, the solution was incubated with trehalase (0.025 U/ml; Sigma-Aldrich, T8778) at 37°C overnight with gentle rotation. For glycogen measurement, the solution was incubated with amyloglucosidase (1 U/ml; Sigma-Aldrich, 10115) at 57°C overnight. The amount of glucose released was measured by glucose assay kit (Sigma-Aldrich, GAGO20). The experiment was performed with 3 biological replicates and observed values were plotted using GraphPad Prism. Statistical significance was calculated using unpaired Student’s *t* test.

### Metabolite extraction and LC-MS/MS analysis

The metabolite extraction and analysis were carried out following protocols described in (Walvekar *et al*, 2019). For each experiment, 10 OD600 cells were used for metabolite extraction. Metabolites were separated on Synergi 4-μm Fusion-RP 80 Å (150 × 4.6 mm) LC column (Phenomenex, 00F-4424-E0) using the Waters Acquity UPLC system. The solvents used for separating sugar phosphatases and trehalose are the following: 5 mM ammonium acetate in water (solvent A) and acetonitrile (solvent B). Solvents used for separating amino acids are 0.1% formic acid in water (solvent A) and 0.1% formic acid in methanol (solvent B). The metabolites were detected using ABSciex QTRAP 6500 mass spectrometer. The data was acquired using Analyst 1.6.2 software (Sciex) and analyzed using MultiQuant version 3.0.1 (Sciex). Analyzed data was plotted using GraphPad Prism.

### ^13^C carbon flux measurements

For measuring carbon flux towards gluconeogenesis, cells were grown in SCD medium for 24hrs and then pulsed with 1% ^13^C2-acetate (Cambridge Isotope Laboratories, CLM-440). Metabolites were extracted after 30 minutes of labelled acetate addition. Samples were analyzed as described earlier. The addition of intensity of all labelled intermediates detected for a metabolite was calculated as total 13C label incorporation. The total label incorporation was normalized with WT values to calculate relative 13C label incorporation. Statistical significance was calculated using paired Student’s *t* test.

### Ethanol estimation assay

The ethanol estimation was carried out as described in (Vengayil *et al*, 2022). Briefly, cells were grown in YPD medium to an OD600 of ∼0.8, cells were centrifuged and 5ml of supernatant was collected in a fresh tube. 1 ml of Tri-n-butyl phosphate (TBP) was added and vortexed for 5 minutes. After centrifugation top layer was transferred to a fresh tube and mixed with an equal volume of potassium dichromate. The mixture was incubated for 10 minutes and absorbance was measured at 595nm. Statistical significance was calculated using unpaired Student’s *t* test.

### Fluorescence microscopy

Cells expressing fluorescently tagged proteins were cultured in YPD medium for 24hrs and then visualized using a microscope (Olympus BX53) with 100 X objective. To stain mitochondria, cells were incubated with Mitotracker Red CMXRos. Far8 was tagged with mNeonGreen for visualizing the Far complex. Sec63-mCherry was used to visualize ER. The experiments were repeated using 3 biological replicates.

### Competition growth-fitness assay

WT cells expressing mCherry and *ppg1Δ1* cells expressing mNeonGreen from chromosomal loci were cultured separately in YPD medium. These cultures were mixed together at an A600 of 0.2 each in YPD medium. Cultures were grown for 24hrs and then subcultured at an A600 of 0.2 in fresh YPD medium. The cells were allowed to grow for 24hrs and were again shifted to a fresh YPD medium, and these transfers were repeated several times. The relative proportion of WT and *ppg1Δ1* cells were estimated by counting the number of mCherry and mNeonGreen positive cells using fluorescence microscopy. The experiment was performed with 3 biological replicates and a minimum of 100 cells were counted for each replicate. As a control, fluorescently labelled WT and *ppg1Δ1* cells were cultured together in YPD medium for 6hrs, and cells were transferred to a fresh YPD medium, and these transfers were repeated. The fractional composition of WT and *ppg1Δ1* cells was calculated and plotted.

### Data visualization and statistical test

All the graphs were plotted and analyzed using GraphPad Prism 6. Two-tailed unpaired Student’s *t* test was used to estimate statistical significance unless otherwise specified. For the flux experiment, the raw intensity values for each replicate were normalized to respective wild-type control and paired Student’s *t* test was used to estimate statistical significance. For the mass spectrometry experiment examining metabolite levels in Far complex mutants, a one-tailed paired Student’s *t* test was used to estimate statistical significance. P values and *n* for corresponding experiments have been specified in figure legends.

## Author contributions

SN and LK contributed in the design and execution of experiments. SN and SL conceptualized the project, analyzed data, and wrote the manuscript.

## Acknowledgements

We acknowledge extensive use of and support from the NCBS-inStem- CCAMP mass spectrometry facility. SL acknowledges a DBT-Wellcome India Alliance Senior Fellowship (IA/S/21/2/505922) and DBT-inStem for support.

## Supplement Figures

**Figure S1:**
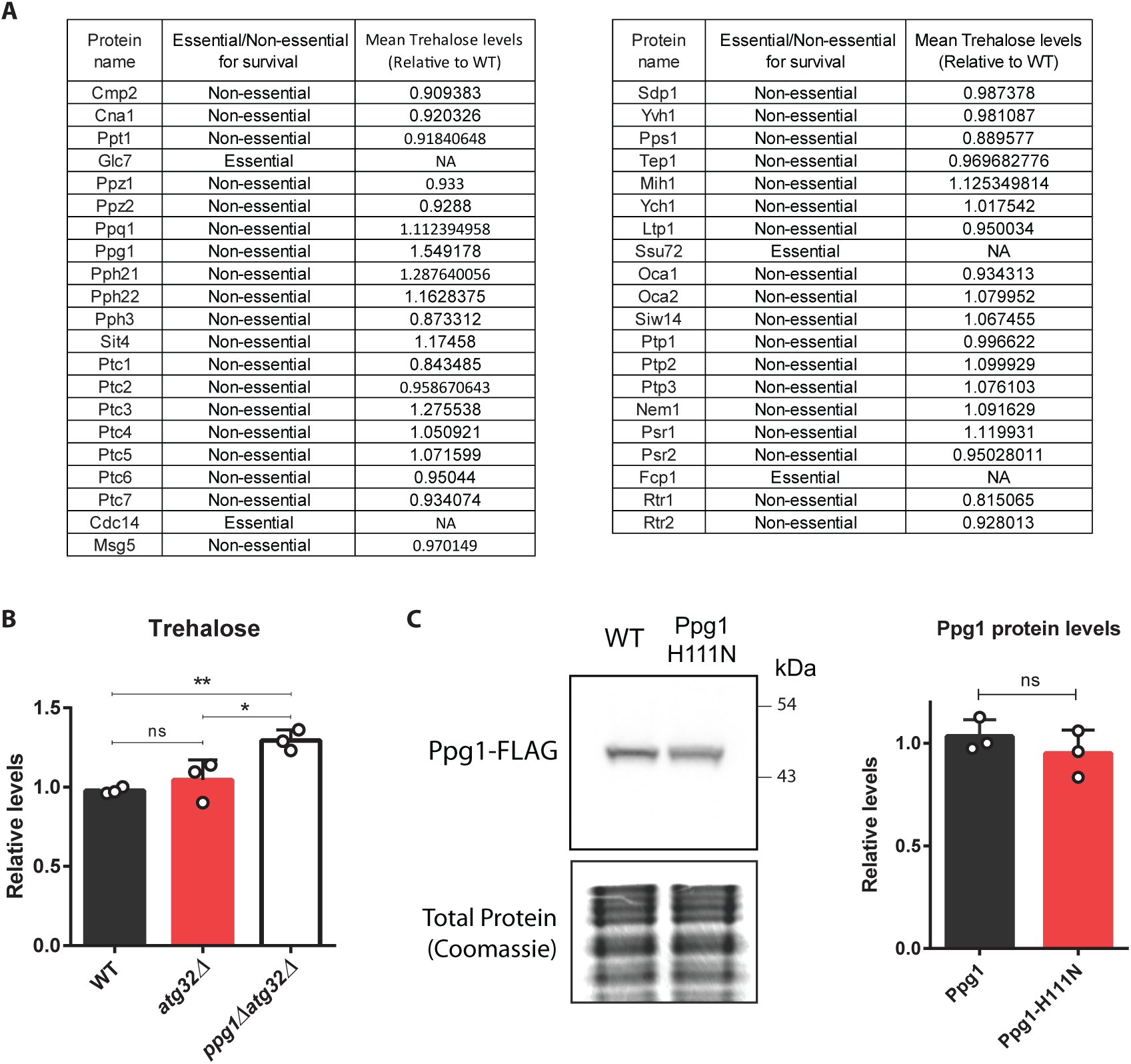
A) A list of protein phosphatases in *S. cerevisiae* and the phosphatase mutants used in the study. Trehalose accumulation was measured after 24hrs of growth in YPD medium. The mean trehalose accumulation was obtained from 2 biological replicates. B) Effect of deletion of Atg32 on trehalose accumulation in WT and *ppg1Δ1* cells. Trehalose accumulation was measured after 24hrs of growth in YPD medium. Data represented as a mean ± SD (n=3). *P < 0.05, **P < 0.01, and ***P< 0.001; n.s., non-significant difference, calculated using unpaired t tests. C) Effect of H111N point mutation on protein levels of Ppg1. The WT and Ppg1H111N cells containing endogenously tagged Ppg1 with 3xFLAG epitope were cultured in YPD medium. Cells were collected after 24hrs of growth and the levels of Ppg1 were measured by western blotting. A portion of the gel was Coomassie stained and used as a loading control. Western blot quantification was done using ImageJ software. A representative image is shown (n=3). Quantification data represented as a mean ± SD (n=3). *P < 0.05, **P < 0.01, and ***P< 0.001; n.s., non-significant difference, calculated using unpaired t tests.

**Figure S2:**
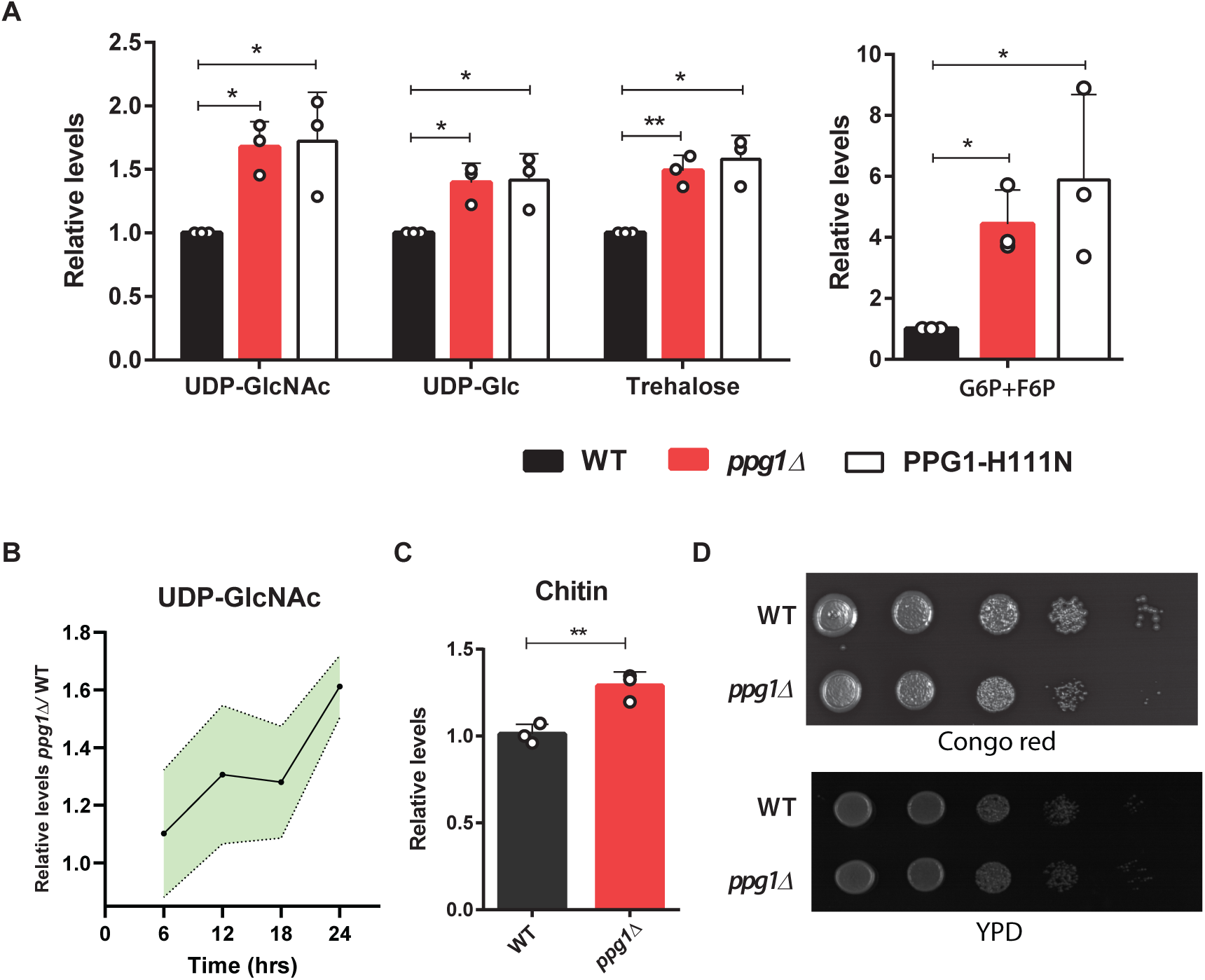
A) Relative steady-state amounts of specific gluconeogenic intermediates, precursors of cell wall and storage carbohydrates, and amino acids in WT, *ppg1Δ1*, and Ppg1H111N cells after 24hrs of growth in YPD medium. Data represented as a mean ± SD (n=3). *P < 0.05, **P < 0.01, and ***P< 0.001; n.s., non-significant difference, calculated using paired t tests. B) Relative steady-state amounts of UDP-GlcNAc from WT and *ppg1Δ1* cells during the course of growth in YPD medium. WT and *ppg1Δ1* cells were grown in YPD medium, and metabolite extraction was carried out. Data represented as a mean ± SD (n=3). C) Relative chitin levels in cell walls of WT and *ppg1Δ1* cells after 24hrs of growth in YPD medium. Data represented as a mean ± SD (n=3). *P < 0.05, **P < 0.01, and ***P< 0.001; n.s., non-significant difference, calculated using unpaired t tests. D) The growth of WT and *ppg1Δ1* cells in presence of Congo red. A serial dilution growth assay was carried out in presence of Congo red using WT and *ppg1Δ1* cells. Congo red was used at a final concentration of 400 μg/ml. The images were taken after 48hrs of growth. A representative image is shown (n=3).

**Figure S3:**
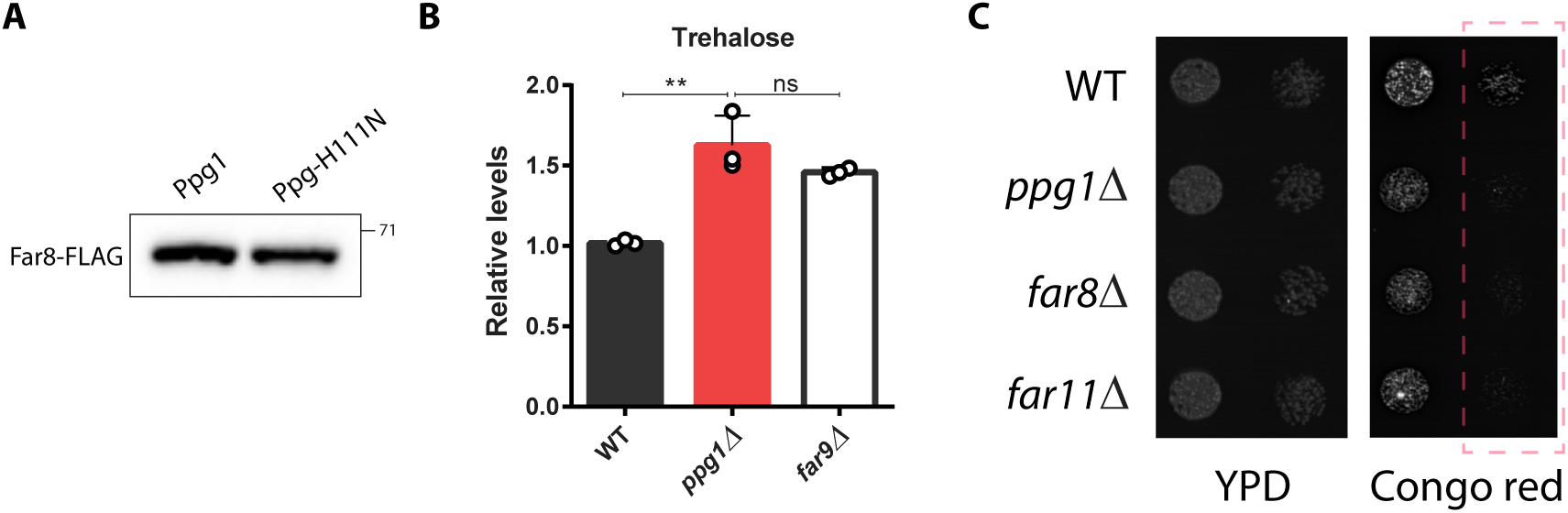
A) Regulation of Far8 post-translational modifications by Ppg1. WT and Ppg1H111N cells containing endogenously tagged Far8 with 3xFLAG epitope were cultured in YPD medium for 24hrs. Far8 mobility was monitored on a 7% SDS-PAGE gel. B) Relative trehalose levels in WT, *ppg1Δ1*, and *far9Δ1* cells after 24hrs of growth in YPD medium. Data represented as a mean ± SD (n=3). *P < 0.05, **P < 0.01, and ***P< 0.001; n.s., non-significant difference, calculated using unpaired t tests. C) The growth of *far8Δ1*, and *far11Δ1* cells in presence of Congo red. A serial dilution growth assay was carried out in presence of Congo red using WT, *ppg1Δ1*, *far8Δ1*, and *far11Δ1* cells. Congo red was used at a final concentration of 400 μg/ml. The images were taken after 48hrs of growth. A representative image is shown (n=3).

**Figure S4:**
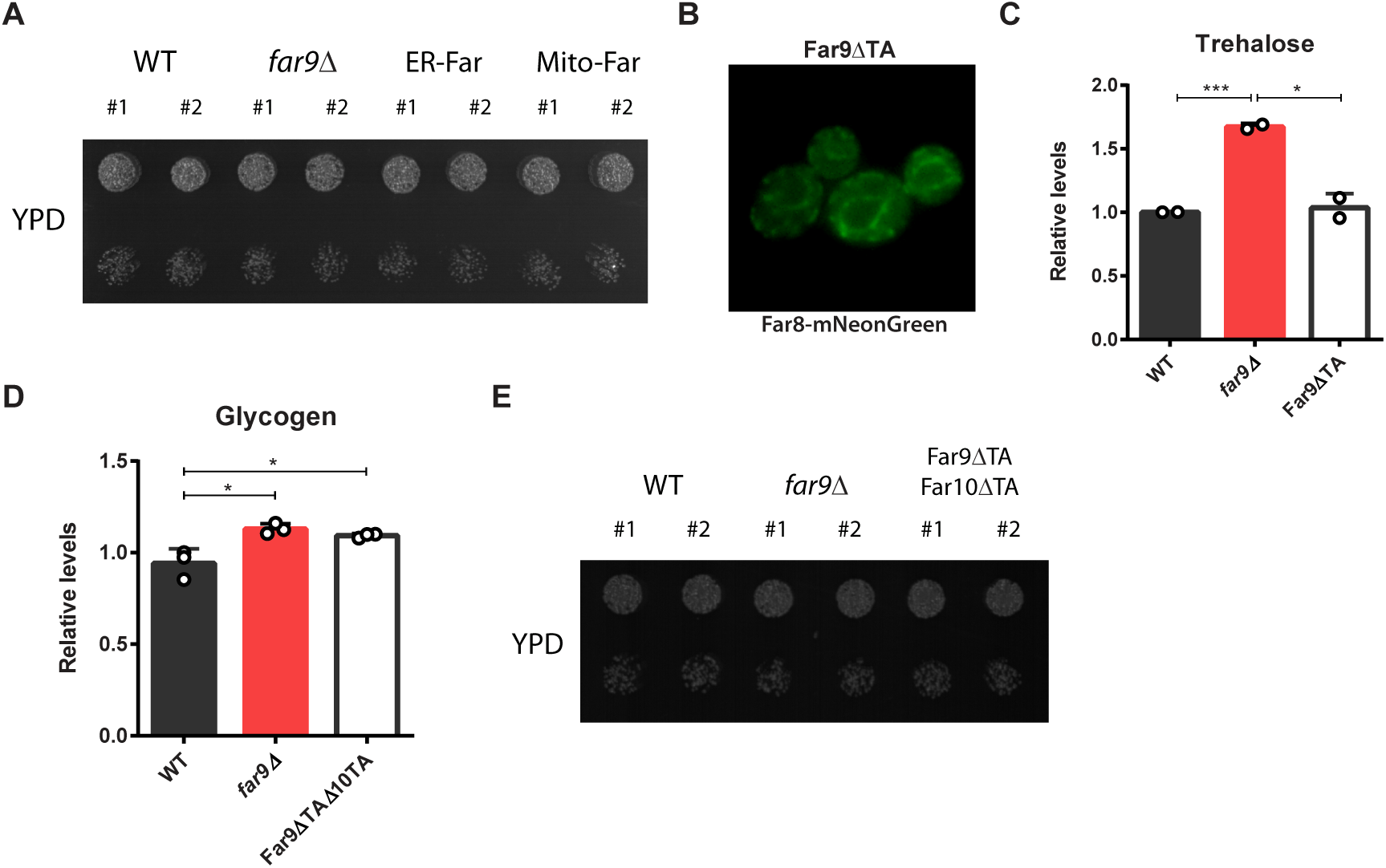
A) The growth of Mito-Far and ER-Far cells in YPD medium. A serial dilution growth assay was carried out using WT, Mito-Far, ER-Far, and *far9Δ1* cells. The images were taken after 24hrs of growth. A representative image is shown (n=2). B) The subcellular localization of Far complex in Far9*⊗*1TA cells. Far8-mNeonGreen was used to visualize the localization of Far complex in these cells. C) Relative trehalose levels in WT, *far9Δ1*, and Far9*⊗*1TA cells after 24hrs of growth in YPD medium. Data represented as a mean ± SD (n=2). *P < 0.05, **P < 0.01, and ***P< 0.001; n.s., non-significant difference, calculated using unpaired t tests. D) Relative glycogen levels in WT, *far9Δ1*, and Far9*⊗*1TA10*⊗*1TA cells after 24hrs of growth in YPD medium. Data represented as a mean ± SD (n=3). *P < 0.05, **P < 0.01, and ***P< 0.001; n.s., non-significant difference, calculated using unpaired t tests. E) The growth of Far9*⊗*1TA10*⊗*1TA cells in YPD medium. A serial dilution growth assay was carried out using WT, *far9Δ1*, and Far9*⊗*1TA10*⊗*1TA cells. The images were taken after 24hrs of growth. A representative image is shown (n=2).

**Figure S5:**
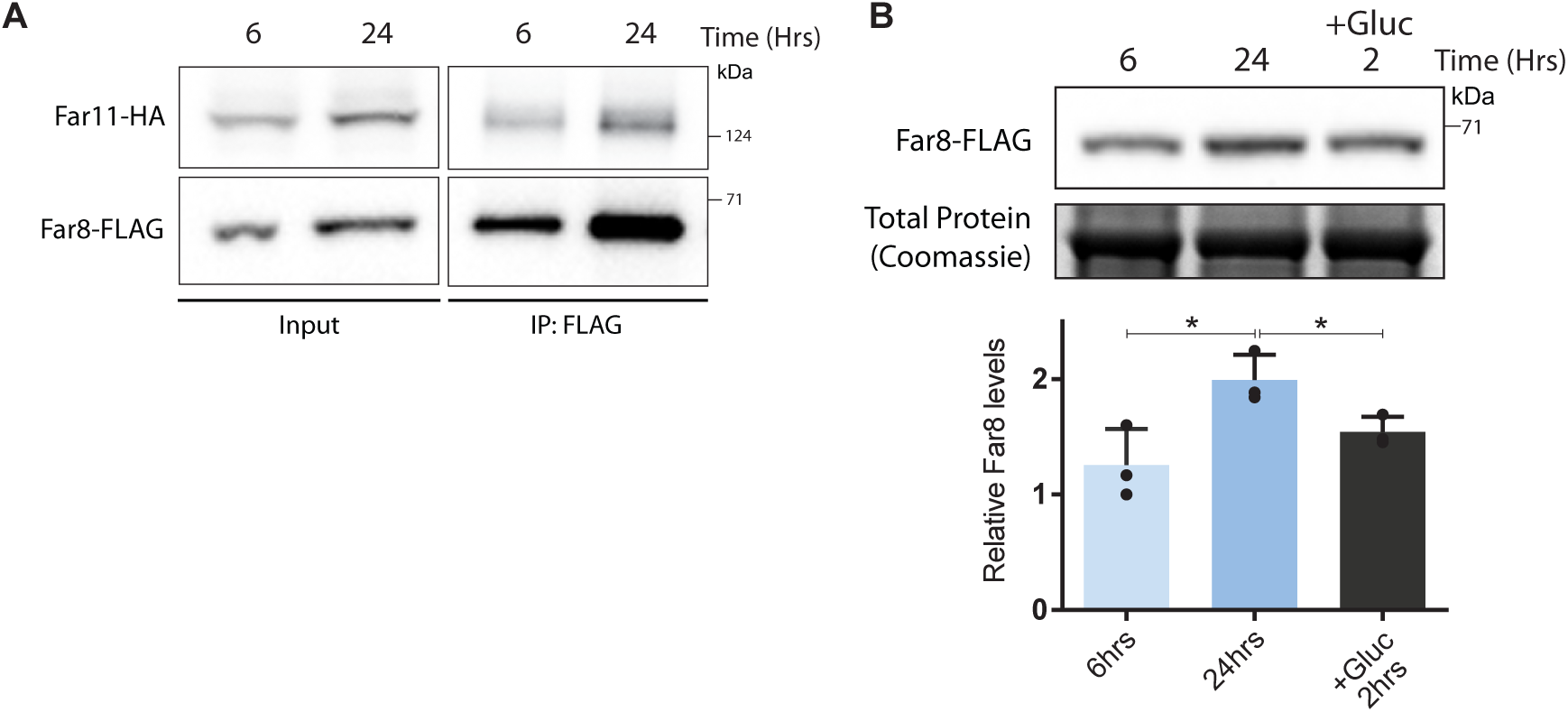
A) Interaction between Far8 and Far11 in different phases of growth. WT cells containing HA-tagged Far11 and FLAG-tagged Far8 were cultured in YPD medium. Cells were collected after 6hrs and 24hrs. Far8-FLAG was immunoprecipitated from these cells, and co-immunopurified Far11-HA was detected. A representative image is shown (n=3). B) Effect of glucose availability on amounts of Far8. Cells expressing FLAG-tagged Far8 were cultured in YPD medium. After 24hrs of growth cells were diluted in the spent medium and 2% glucose was added. Cells were collected after 2hrs and levels of Far8 were measured by western blotting. A portion of the gel was Coomassie stained and used as a loading control. Western band quantification was done using ImageJ software. A representative image is shown (n=3). *P < 0.05, **P < 0.01, and ***P< 0.001; n.s., non-significant difference, calculated using unpaired t tests.

**Figure S6:**
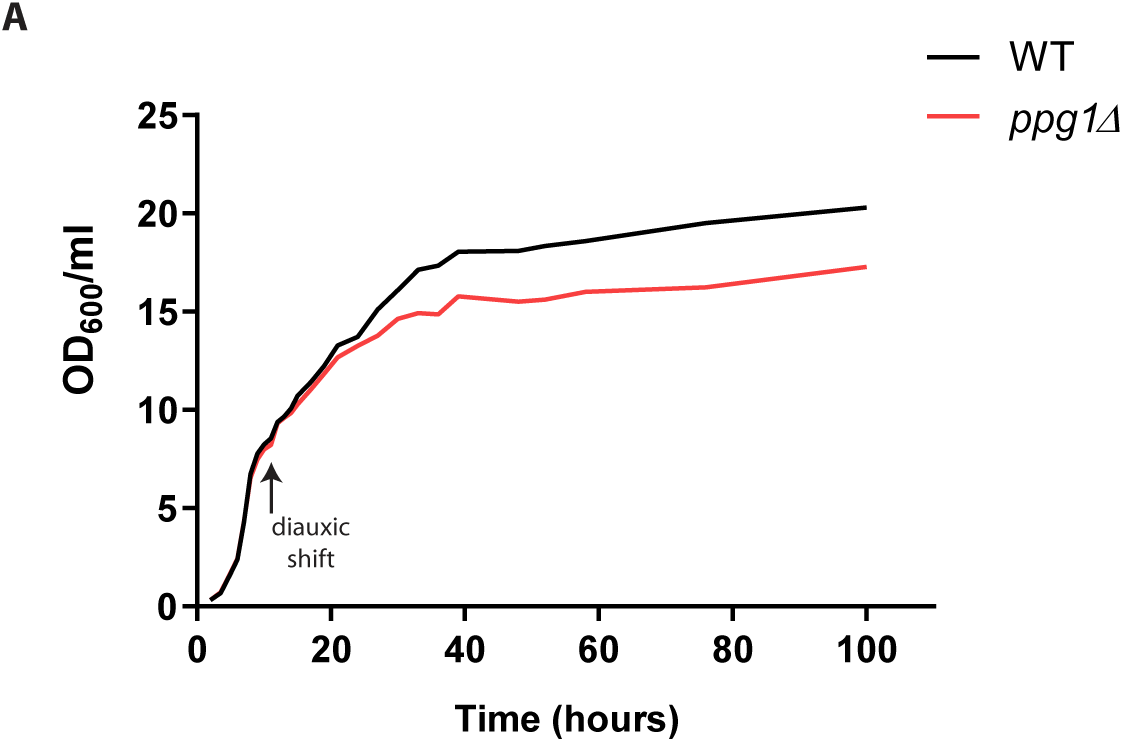
A) Growth dynamics of *ppg1Δ1* cells in YPD medium. The cultures of WT and *ppg1Δ1* were started at OD600 of 0.2 in YPD medium and the growth was monitored.

